# GreA and GreB enhance *Escherichia coli* RNA polymerase transcription rate in a reconstituted transcription-translation system

**DOI:** 10.1101/024604

**Authors:** Lea L. de Maddalena, Henrike Niederholtmeyer, Matti Turtola, Zoe N. Swank, Georgiy A. Belogurov, Sebastian J. Maerkl

## Abstract

Cell-free environments are becoming viable alternatives for implementing biological networks in synthetic biology. The reconstituted cell-free expression system (PURE) allows characterization of genetic networks under defined conditions but its applicability to native bacterial promoters and endogenous genetic networks is limited due to the poor transcription rate of *Escherichia coli* RNA polymerase in this minimal system. We found that addition of transcription elongation factors GreA and GreB to the PURE system increased transcription rates of *E. coli* RNA polymerase from sigma factor 70 promoters up to 6-fold and enhanced the performance of a genetic network. Furthermore, we reconstituted activation of natural *E. coli* promoters controlling flagella biosynthesis by the transcriptional activator FlhDC and sigma factor 28. Addition of GreA/GreB to the PURE system allows efficient expression from natural and synthetic *E. coli* promoters and characterization of their regulation in minimal and defined reaction conditions making the PURE system more broadly applicable to study genetic networks and bottom-up synthetic biology.

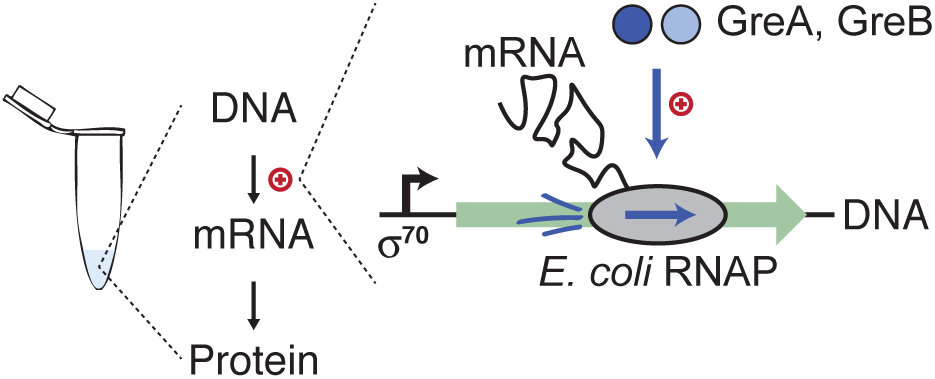
Graphical abstract.

## Introduction

Diverse biological processes can be reconstituted and studied *in vitro* using purified proteins or lysates. This approach has facilitated fundamental discoveries in molecular biology and biochemistry, such as DNA replication^1^ and translation of the genetic code^2^. The cell-free approach allows construction of systems that would be difficult or impossible to develop *in vivo* and to perform measurements and experiments that are difficult to conduct in cells. Apart from basic research on biological networks, applications of *in vitro* systems thus far have include biosensors and molecular synthesis ^3–5^. *E. coli* cell-lysate transcription and translation (TX-TL) systems have become popular to engineer and study genetic networks of increasing complexity^6–10^. Cell-lysate-based TX-TL systems produce high protein yields and allow transcription from native *E. coli* promoters^11^. However, lysates have the disadvantage that they contain almost all the proteins and macromolecules present in the cytoplasm at the moment of lysis. For bottom-up synthetic biology and reconstitution studies this is not ideal because the lysate still contains many unknown components and uncontrollable variables.

An appealing transcription and translation system that is commercially available as PURExpress (PURE), is reconstituted from purified components and allows experiments under minimal and defined conditions^12^. This reconstituted system contains T7 RNA polymerase (RNAP), purified ribosomes, all necessary translation factors from *E. coli*, tRNAs, enzymes for tRNA aminoacylation and energy regeneration, creatine phosphate as an energy source, and nucleotides and amino acids as precursors. Various genetic networks can be implemented in this system, which to date mostly relied on single subunit phage RNA polymerases for transcription^13^.

Transcription by the multisubunit *E. coli* RNA polymerase (EcRNAP) has been reconstituted in the PURE system and can be implemented by either adding the purified holoenzyme to the reaction mix or by co-expressing its subunits^14,15^. While transcription rates of the EcRNAP in PURE depend on the concentrations of DNA template and EcRNAP, they are generally considerably lower than for phage polymerases. For example, we observed roughly an order of magnitude lower mRNA concentrations synthesized from a consensus sequence sigma factor 70 (σ70) promoter by EcRNAP than by phage T3 RNAP^13^ under similar conditions. *In vivo* elongation rates of EcRNAP range between 28 and 89nt/s^16^ and are comparable to the values reported for T7 RNAP *in vitro*.^17,18^ Transcription elongation factors affect EcRNAP transcription elongation rate either by sensitizing or suppressing RNAP pausing. Elongating RNAP frequently backtracks along the DNA forming a transcriptionally inactive state. Transcription elongation factors GreA and GreB bind to backtracked RNAP and catalyze the endonucleolytic cleavage of nascent RNA within the RNAP active site allowing transcription to continue^19^. GreA resolves smaller backtracking events by cleaving 2-3 nt from the 3’end of the RNA, whereas GreB can also rescue longer backtracked complexes^20^. GreA and GreB are known to increase transcription elongation rate and stimulate promoter escape in a subset of promoters^19–22^, but GreA and GreB transcription elongation factors are not present in the PURE system.

Here we show that *E. coli* transcription elongation factors GreA and GreB enhance EcRNAP transcription rates in the PURE system up to 6-fold to reach the rate of T7 RNAP transcription in the system. We go on to show that an increase in transcription rates can be observed for several different synthetic and natural σ70 *E. coli* promoters. Furthermore, we used the enhanced system to study natural *E.coli* promoters involved in flagella biosynthesis and their activation by two different transcriptional activators *in vitro*, under defined conditions.

## Results and discussion

EcRNAP can be added to the PURE system to allow transcription of DNA templates carrying *E. coli* promoters^14,15^ but mRNA synthesis and subsequent protein production is more efficient using phage polymerases such as T7 or T3 RNAP^13^. In bacterial cells multiple proteins can increase RNAP activity, which are not present in the minimal PURE system. The transcription elongation factors GreA and GreB from *E. coli* increase overall transcription elongation rates and stimulate promoter escape in a subset of promoters by re-activating backtracked elongation complexes^19–22^. We added transcription elongation factors GreA and GreB to a PURE reaction containing EcRNAP with σ70 (holoenzyme) and a DNA template expressing EGFP under control of a consensus sequence *E. coli* σ70 promoter. We measured concentrations of full-length mRNA using fluorescent FRET probes that bind to a target region at the 3’ end of the mRNA^23^ (Fig. 1A) and found that in the presence of GreA and GreB mRNA concentrations increased faster and to higher concentrations than without the elongation factors (Figure 1). The transcription rate increase mediated by GreA and GreB followed hyperbolic kinetics and plateaued at concentrations above 5 μM for both GreA and GreB. At the plateau GreA and GreB increased transcription rates about 3-fold and final EGFP protein synthesized about 2-fold for the DNA template concentration tested (Fig. 1B, C). When GreA and GreB were added in combination, we did not observe a significant synergistic effect on mRNA or protein synthesis (Supplementary Fig. 1). We nonetheless used both proteins in an enhanced PURE (ePURE) reaction containing 10 μM of GreA and 10 μM of GreB in addition to the *E. coli* RNAP holoenzyme (0.2 μM EcRNAP, 1 μM σ70) for all subsequent experiments. Apart from these protein additions we did not further modify the commercial PURE system (see Materials and Methods). During a TX-TL reaction in ePURE we observed up to 10-fold higher mRNA concentrations, which also translated to an increased EGFP synthesis (Fig. 1D, E).

**Fig. 1:**
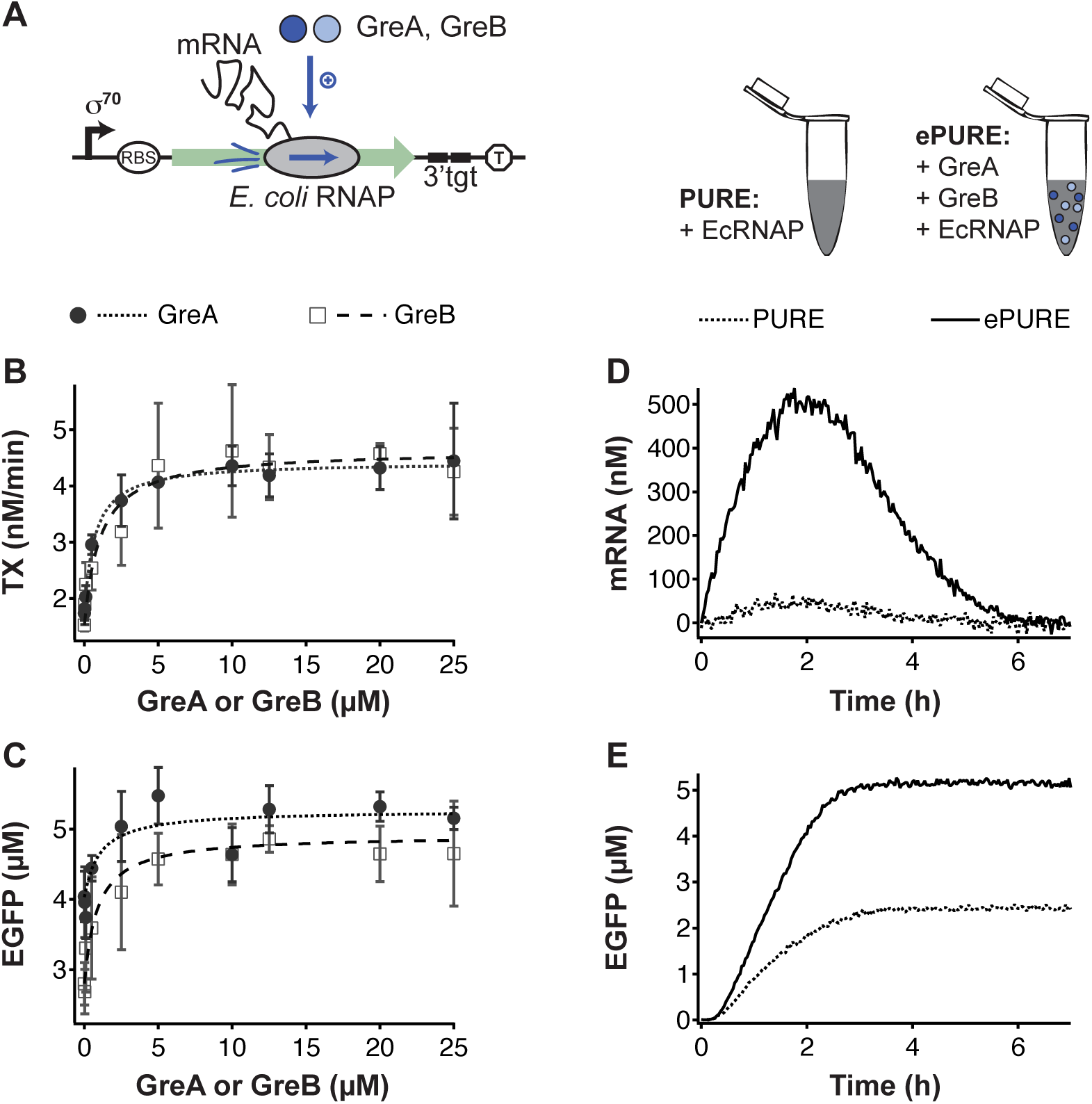
GreA and GreB transcription elongation factors increase EcRNAP transcription rates in a PURE TX-TL reaction. **(A)** GreA and GreB proteins enhance transcription of a DNA templates by EcRNAP in a PURE TX-TL reaction. The DNA template encoded EGFP under control of a consensus sequence σ70 promoter and a strong ribosomal binding site (RBS). Upstream of a transcriptional terminator it carried target region (3’ tgt) to measure the concentration of full-length mRNA using binary FRET probes. Transcription rates **(B)** and final EGFP concentrations **(C)** increase with increasing GreA and GreB concentrations following hyperbolic kinetics. Lines represent fits to the Michaelis-Menten equation. All GreA and GreB concentrations were tested at least in duplicate. **(D, E)** Measurements of mRNA and EGFP concentrations over time during a TX-TL reaction in PURE and ePURE. Transcription rates and final EGFP concentrations were determined from traces like these as described in the Materials and Methods section. DNA templates carried the σ^70^tet promoter and was used at a concentration of 8 nM.

By testing various DNA template concentrations we observed an up to 6-fold increase of transcription rate in ePURE compared to a PURE reaction supplemented with *E. coli* RNAP holoenzyme alone (Fig. 2A). Increased mRNA synthesis led to 3-fold higher final EGFP levels, and the advantage of using the ePURE reaction was strongest for lower DNA template concentrations (Fig. 2B). In the ePURE system we observed comparable mRNA and EGFP synthesis from an *E. coli* σ70 promoter and a T7 RNAP promoter (Fig. 2). The ePURE improvement should thus facilitate using *E. coli* RNAP for transcription under defined conditions.

**Fig. 2:**
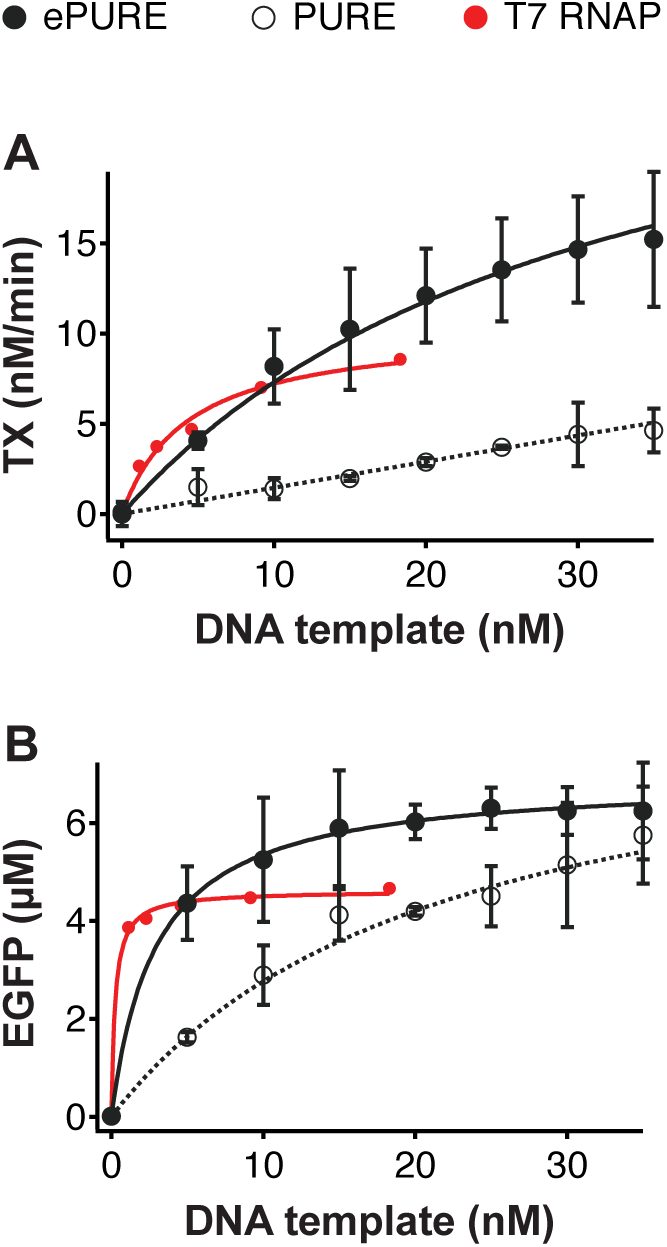
Enhanced transcription in the ePURE system as a function of DNA template concentration. Effect of varying DNA template concentrations on transcription rates **(A)** and final EGFP concentration **(B)**. The DNA template carried the σ^70^tet promoter and DNA concentrations were tested in two independent experiments. Data for comparison to the T7 RNAP was previously collected in a PURE reaction without EcRNAP and GreA and GreB proteins^23^. Lines represent fits to the Michaelis-Menten equation.

We next asked whether GreA and GreB can improve transcriptional elongation rates for genes other than EGFP. In order to test this, we measured synthesis of full-length mRNA from two additional genes: *asr* (309 bp), and *chiP* (1407 bp) which are roughly half and twice as long as the EGFP gene (717 bp), respectively. Transcription rates of both genes were increased in the ePURE system in the presence of GreA and GreB (Supplementary Fig. 2). In the non-optimized PURE system transcription rates were below the detection limit for the longer gene and increased to measurable levels in the presence of GreA and GreB. The transcription rate of the shorter gene increased 8-fold (Supplementary Fig. 2).

In order to determine if the ePURE system also increases the transcription rate for promoters other than the strong σ70tet promoter we characterized in Figures 1 and 2, we tested the system on 16 additional synthetic and natural σ70 promoters (Fig. 3). The panel was composed of nine constitutive promoters from the registry of standard biological parts (http://parts.igem.org), the BBa_J231xx-series, which are well-characterized *in vivo* and *in vitro*,^24–26^, two constitutive promoters *proC* and *proD* ^27^, several natural repressible promoters (promoter of the *trp* operon, *lac* UV5 promoter, the phage λ_PR_ promoter), and three synthetic repressible promoters (P_tet_ ^28^, and the σ70 consensus sequence promoters P_*σ*70_lac and P_σ70_tet ^13^). In the ePURE system EGFP synthesis increased for 14 of the 17 promoters we characterized (Fig. 3A). Our results on the constitutive J231xx-promoters compare well with relative promoter strengths measured in lysate-based TX-TL reactions^25,26^. We found that the presence of GreA and GreB in the ePURE system enhanced expression particularly for strong promoters. These results suggest that at least for strong promoters transcriptional pausing during elongation severely limited transcription by EcRNAP in the PURE system. However, we cannot rule out effects on initiation of transcription as it has been shown that GreA and GreB can influence promoter escape differently for different promoters^22^. By increasing the range of synthesis rates that can be attained with *E. coli* promoters the ePURE system will be useful in implementing and characterizing genetic networks based on native *E. coli* promoters.

**Fig. 3:**
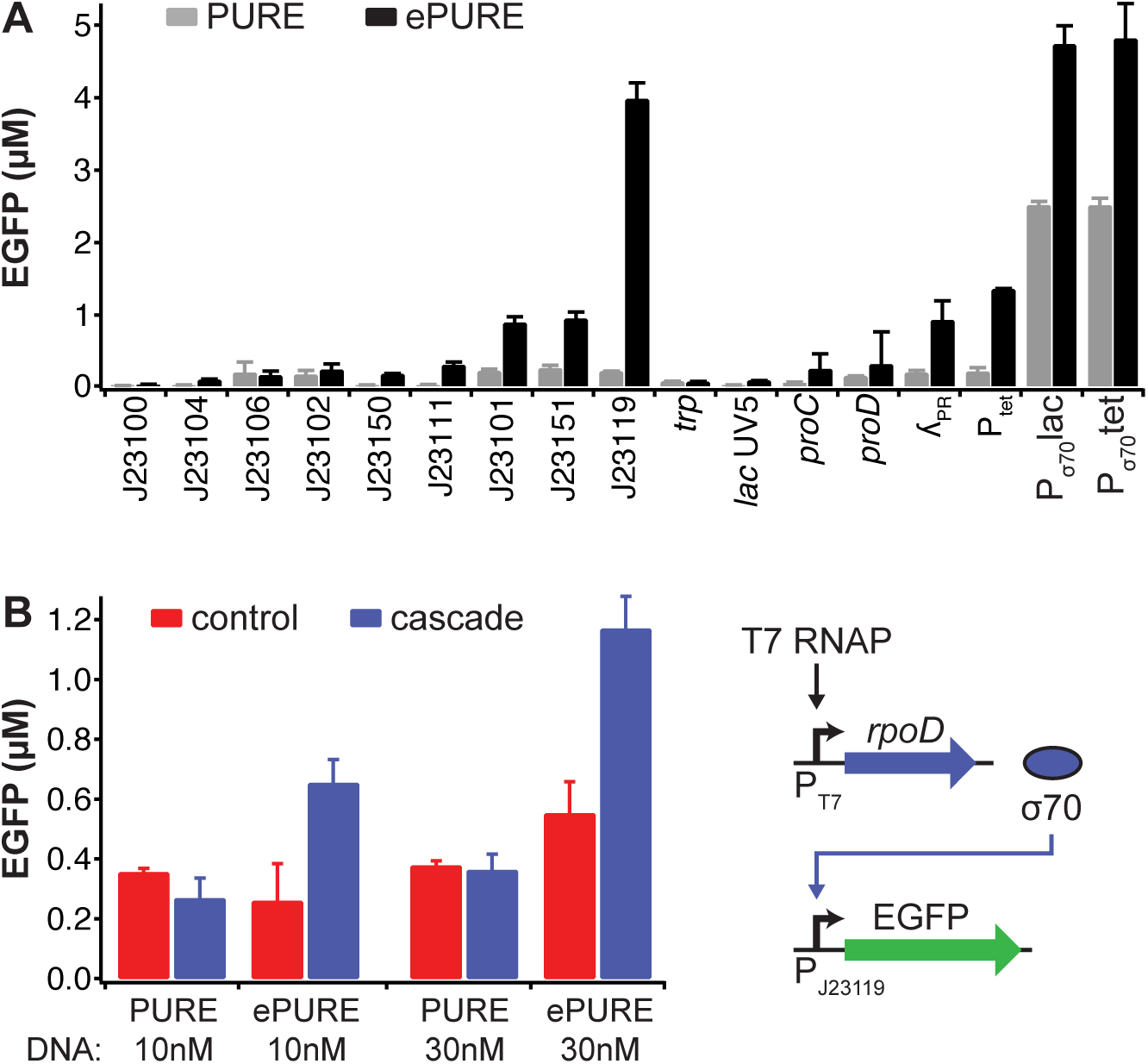
ePURE enhances synthesis from different σ70 EcRNAP promoters and performance of a genetic cascade. (A) Comparison of final EGFP produced from different σ70 EcRNAP promoters in the PURE and the ePURE system. DNA templates were at a concentration of 8 nM and encoded EGFP. (B) Comparison of the performance of a synthetic 2-gene circuit in the PURE and the ePURE system at low and high DNA template concentrations. Shown are endpoint EGFP concentrations. For the full cascade, both DNA templates of the circuit shown on the right were used at 10 nM and 30 nM respectively. For the control reactions, the *rpoD* template was omitted. To observe activation of the J23119 promoter by σ70, PURE and ePURE in (B) were assembled with only the core EcRNAP. Values are averages of two independent experiments with error bars showing the standard deviation.

As an example of this we assembled a simple synthetic 2-gene cascade that consisted of T7 RNAP transcribing the *rpoD* gene to synthesize σ70, which then, together with EcRNAP core enzyme, activated EGFP expression (Fig. 3B). In order to study the effect of DNA template concentration on the performance of the network, we additionally analyzed the circuit at a low and a high DNA template concentration of 10 and 30 nM, respectively (Fig. 2B). The PURE system failed to run the genetic cascade at either DNA concentration and yielded output comparable to the negative control lacking the *rpoD* template. The ePURE system on the other hand yielded increased expression of the reporter, which was further boosted at higher DNA concentrations. This demonstrates that addition of GreA and GreB enables the implementation of genetic networks involving EcRNAP. In fact, in this case ePURE was necessary for the cascade to function, and simply increasing DNA template concentrations was not sufficient to improve performance of the network in the PURE system.

We next tested whether the ePURE system would allow us to study an endogenous bacterial genetic network and chose to analyze transcriptional regulation of native *E. coli* flagellar promoters. Two main regulators, the FlhDC transcriptional activator and FliA, the flagellar sigma factor, σ28, are thought to activate the genes in a tightly controlled temporal order^29,30^. FlhDC is known as the master regulator and activates σ70-dependent transcription from class 2 promoters. One of the genes FlhDC activates is *fliA* coding for σ28, which then activates itself and other genes in a positive autoregulation^31^. Many of the more than 50 genes in the flagellar regulon, which are divided into at least 17 operons, are transcribed from multiple promoters, and can be activated by both FlhDC and σ28^29,32^. Their regulation has been studied extensively *in vivo* using genetics and promoter fusions^29,30,32^. In a complementary approach, this complex regulatory system can also be studied *outside of cells*, under reaction conditions that eliminate unknown factors. Activation of several flagellar class 2 and class 3 promoters by FlhDC and σ28 was shown by *in vitro* transcription experiments^31,33,34^ and binding of the FlhDC activator to putative promoters was demonstrated by *in vitro* binding assays^35^.

We analyzed eight flagellar promoters coupled to an EGFP reporter in the ePURE system and show their activation by FlhDC and σ28 in the defined TX-TL system (Fig. 4A). We based our expectations on the EcoCyc *E. coli* database^36^, which contains information on experimentally known and bioinformatically predicted transcriptional activation. To synthesize our DNA templates we fused 150 to 250 bp long promoter regions that contained the annotated σ70 and σ28 promoters as well as FlhDC binding sites to identical, strong ribosomal binding sites followed by the EGFP reporter gene. To test their activation, we separately pre-synthesized the FlhDC and the σ28 activators and then added these to an ePURE reaction containing GreA, GreB, EcRNAP, and a DNA template with the respective flagellar promoters.

**Fig. 4:**
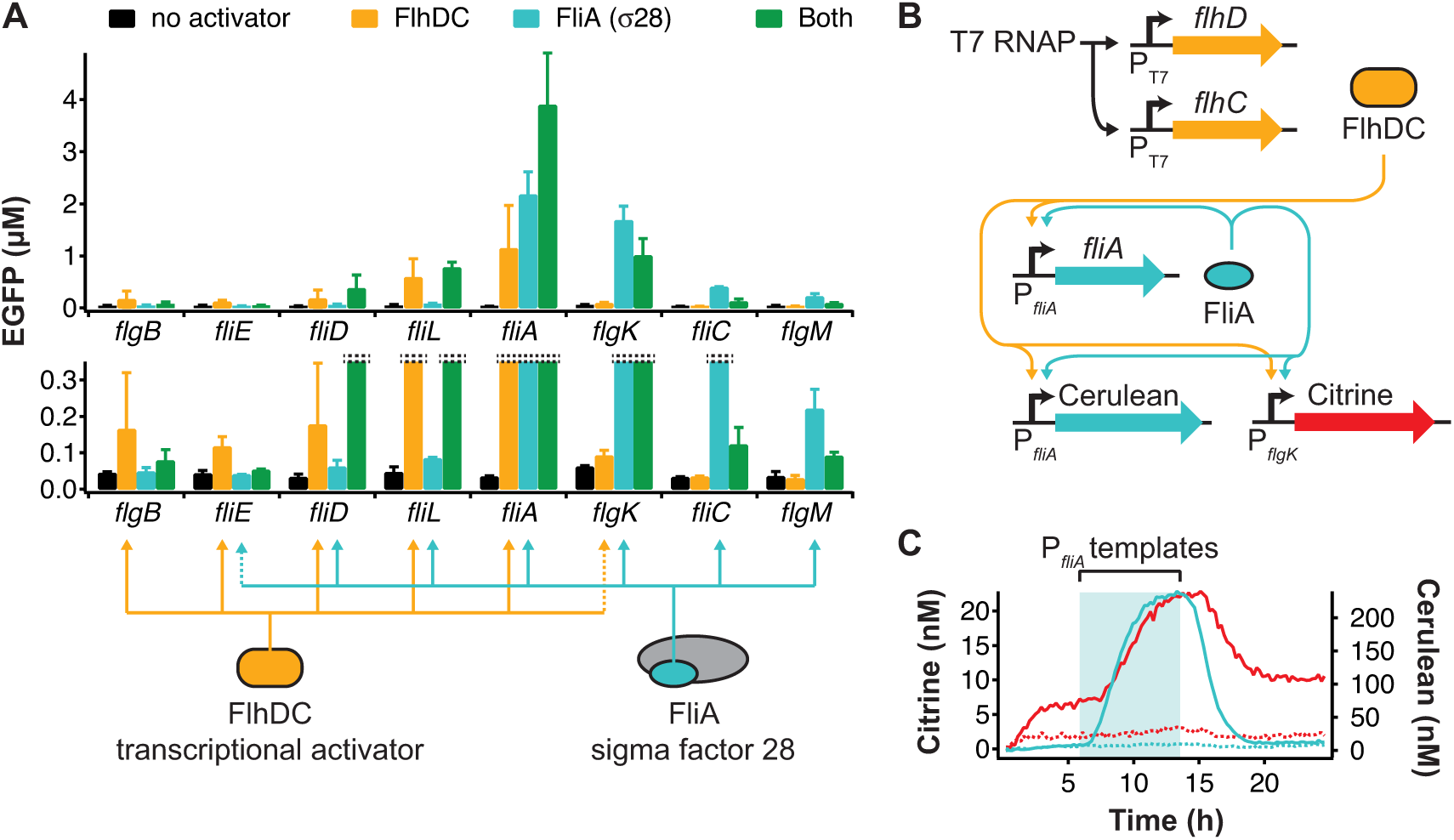
*E. coli* flagellar promoter activation by FlhDC and σ28 factor in ePURE. **(A)** Addition of pre-synthesized FlhDC and σ28 to an ePURE reaction activated EGFP expression from eight native *E. coli* promoters involved in flagella synthesis. The activators were added separately or in combination. The activation pattern expected from annotations on EcoCyc is shown beneath the experimental results. Dotted lines represent deviations of our results from the expectation and are discussed in the text. All promoter-EGFP templates were used at 6.5nM. Values are averages of two independent experiments with error bars showing the standard deviation. **(B)** A 5-gene genetic network built from FlhDC and σ28 activators and two native flagellar promoters control expression of Citrine and Cerulean reporters. **(C)** The flagella gene network was characterized in a microfluidic nano-reactor device in a continuous ePURE reaction. The P_*fliA*_-*fliA* and P_*fliA*_-Cerulean templates were added transiently during the shaded region of the plot. Solid lines are Cerulean (cyan) and Citrine (red) concentrations for the full network, dotted lines represent a control experiment omitting the P_T7_-*flhD* and P_T7_-*flhC* templates. The figure shows a representative result of two experiments.

All eight promoters tested showed no detectable activity in the absence of FlhDC and σ28. When the reaction contained either of the two activators, we observed the expected activation pattern with widely differing promoter strengths (Fig. 4A). Both activators in combination generally did not improve expression compared to only one activator. Most of the time the presence of both activators even led to decreased expression. We attribute this effect to competition between both activators for binding to DNA^31^ and to the RNAP core enzyme, which binds σ28 with a higher affinity than σ70^34^. We hypothesize that concentrations of EcRNAP and activators might be different in cells than in our *in vitro* assay, which could explain why the promoters did not show additive activation as observed *in vivo*^32^. When considering activation by a single activator, two out of eight promoters, *fliE* and *flgK*, deviated from the annotated regulation pattern in EcoCyc. The *fliE* promoter was predicted to be activated by both σ28^37^ and FlhDC^35^ but we only detected low activation by FlhDC. For the *flgK* promoter only activation by σ28 was annotated on EcoCyc and previously shown^34^. We observed strong activation by σ28 but also a low but significant activation by FlhDC. A computational search for FlhDC did not identify a FlhDC binding site upstream of the *flgK* gene in *E. coli*.^35^ In *Salmonella typhimurium*, however, transcription of *flgK* is activated by both σ70/FlhDC and σ28^38^. Additionally, our study provides experimental evidence for activation of several promoters by FlhDC and σ28, which have previously only been predicted computationally, such as activation of *flgB, fliE* and *fliD* by FlhDC^35^ and *flgM* by σ28^39^. Thus, our analysis of flagellar promoters in defined conditions demonstrates that native gene activation mechanisms can be studied using the ePURE system. The finding that FlhDC is a strong transcriptional activator for a number of different promoters should furthermore be useful for the assembly of synthetic genetic networks.

Using the genes for FlhDC and σ28 and two promoters that showed activation by both activators, we built a synthetic gene network (Fig. 4B), which was implemented and characterized in a microfluidic nano-reactor device^13^. T7 RNAP transcribes the genes coding for FlhD and FlhC. FlhDC then activates σ70 *E. coli* RNAP to express the Citrine reporter, which we placed under control of the *flgK* promoter. A control reaction, which did not contain the FlhD and FlhC DNA templates, again demonstrated activation of the *flgK* promoter by the FlhDC activator (Fig. 4C). Transiently we added DNA templates carrying the *fliA* gene (σ28) and a Cerulean reporter gene, both under control of the *fliA* promoter, which leads to positive autoregulation of σ28. σ28 further activates both reporters leading to a fluorescence increase of Citrine and Cerulean in the presence of the *fliA* and Cerulean templates (Fig. 4C). This 5-gene network demonstrates that complex genetic networks dependent on the *E. coli* RNAP and native *E. coli* promoters can be assembled in the ePURE system.

*E. coli* promoters offer a wide and well-characterized repertoire of promoter-regulator pairs and *E. coli* promoters are also highly modular^40^ making transcription by *E. coli* RNAP interesting for *in vitro* synthetic biology^11^. While lysate-based TX-TL systems can be prepared with high activities of the EcRNAP^11,41^, we have found activity of the EcRNAP in the reconstituted PURE system to be too low for many applications. PURE is a minimal system that can be rationally improved by a bottom-up approach. Here we enhanced transcription in the PURE system by adding purified transcription elongation factors GreA and GreB, which indicates that transcriptional elongation was limiting performance of EcRNAP in the PURE system. Addition of GreA and GreB increased EcRNAP transcription rates from a strong σ70 promoter to levels observed with phage T7 RNAP and significantly increased transcription from a number of synthetic *E. coli* promoters of different strengths. We used the ePURE system to characterize activation of native *E. coli* flagellar promoters by the transcriptional activators FlhDC and σ28, demonstrating that the ePURE system is useful for the characterization of genetic regulation under defined conditions. Inclusion of GreA and GreB proteins should furthermore be useful to increase protein yields in the PURE system when it is desirable to use *E. coli* promoters.

## Materials and Methods

### DNA template preparation

Linear DNA templates were produced by two-step PCR as described previously^13,23^ using the primers listed in Table S1. Flagella promoters were PCR amplified from *E. coli* BL21(DE3) genomic DNA and replaced the 5’extension primer during two-step PCR. All linear DNA templates prepared for this study are listed in Table S2.

### Preparation of GreA, GreB and EcRNAP holoenzyme

EcRNAP subunits (expression plasmid pVS10) were co-expressed in *E. coli* Xjb(DE3) cells (Zymo Research, Irvine, CA, USA) and the α_2_ββ′ω assembly (β′ subunit contained C-terminal His6-tag) was purified by a combination of immobilized-metal affinity, heparin and anion exchange chromatographies as described^42^. *E. coli* σ70 protein containing N-terminal His6-tag (expression plasmid pET28-σ70^43^) was expressed in *E. coli* Xjb(DE3) cells and purified by a combination of immobilized-metal affinity, heparin and anion exchange chromatographies as described for EcRNAP except that the lysis and the wash buffers during immobilized-metal affinity chromatography contained 1 M NaCl. *E. coli* GreA and GreB proteins containing C-terminal His_6_-tags (expression plasmids pIA578 and pIA577, respectively) were purified by immobilized-metal affinity chromatography followed by gel filtration as described^44^. All proteins were dialyzed against the storage buffer (20 mM Tris-HCl pH 7.9, 150 mM NaCl (1M NaCl for GreA and GreB), 0.1 mM EDTA, 50% glycerol, 0.1 mM DTT) and stored at -20°C. A stock solution of 50x holoenzyme (1:5 *E. coli* core RNAP : σ70 factor at 10 μM and 50 μM) was prepared by incubating the proteins in storage buffer at 30°C for 20 min, then the stock was stored at -20°C until use.

### Batch reaction setup and measurements

TX-TL reactions were performed in the PURExpress *In Vitro* Protein Synthesis kit (New England Biolabs) supplemented with Protector RNase Inhibitor (Roche), 1 μM Cy3 and Cy5 binary probes^23^, 200 nM *E. coli* core RNAP and 1 μM σ70 factor. The ePURE system additionally contained 10 μM of each GreA and GreB transcription elongation factors. Platereader batch TX-TL reactions, and measurement of the mRNA concentration were performed as previously described^23^. The initial mRNA synthesis, or transcription rate (TX), was determined by fitting the mRNA concentration (m) of the first 40 min of the reaction to:

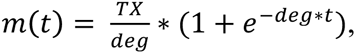

where *t* is time and *deg* signifies the mRNA degradation rate (fixed at 0.0085 min^−1^)^23^. The final EGFP concentration was determined at the plateau of the protein synthesis reaction.

To test activation of flagellar promoters by FlhDC and σ28, we pre-synthesized the activators from T7 RNAP templates in a standard PURE reaction without EcRNAP and Gre proteins. FlhDC was produced by combining *flhD* and *flhC* templates at 10nM each, and σ28 was produced from 10 nM fliA template for 100 min at 37°C. The activators then were stored in aliquots at -80°C until use. These were prepared by combining the FlhDC and σ28 pre-synthesis reactions 1:1 for testing activation by both activators and by diluting 1:1 with Tris buffer for testing of each activator separately.

### Flagellar gene network in a nano-reactor device

We assembled the genetic network from 5 individual DNA templates. Final concentrations were 1 nM for *P*_T7_-*flhD*, 2.5 nM for P_*flgK*_-Citrine, and *P*_T7_-*flhC*, and 2 nM for P_*fliA*_-*fliA* and P*fliA*-Cerulean. The templates P_*fliA*_-*fliA* and P_*fliA*_-Cerulean were only added transiently. The microfluidic chip was prepared and used as described^13^ using a dilution time, t_d_, of 39.6 ±0.4 min. Citrine and Cerulean concentrations were determined from a calibration with purified proteins^10^.

## Supporting Information

Supporting figures S1 and S1. Supporting tables S1 and S2.

## Author Information

Author contributions: LLM and HN contributed equally to this work

## Acknowledgements

The authors would like to thank Irina Artsimovitch (Ohio State University) for providing expression plasmids for EcRNAP, GreA and GreB.

